# Environmental conditions and agronomic practices induce consistent global changes in DNA methylation patterns in grapevine (*Vitis vinifera* cv Shiraz)

**DOI:** 10.1101/127977

**Authors:** Huahan Xie, Moumouni Konate, Na Sai, Kiflu Gebremicael Tesfamicael, Timothy Cavagnaro, Matthew Gilliham, James Breen, Andrew Metcalfe, Roberta DeBei, Cassandra Collins, Carlos Marcelino Rodriguez Lopez

## Abstract

Fruit attributes that affect wine quality are thought to be largely driven by the interaction of grapevine’s genetic characteristics with environmental factors (i.e. climate, soil and topography) and vineyard management. All these variables, in conjunction with the wine making process, give a wine its distinctive character. Understanding how grapevines perceive and adapt to a changing environment will provide us with an insight into how to better manage crop quality. Mounting evidence suggests that epigenetic mechanisms are a key interface between the environment and the genotype that ultimately affect the plant’s phenotype. Moreover, it is now widely accepted that epigenetic mechanisms are a source of useful variability during crop varietal selection that could affect crop performance. While the contribution of DNA methylation to plant performance has been extensively studied in other major crops, very little work has been done in grapevine. Here we used Methylation Sensitive Amplified Polymorphisms to obtain global patterns of DNA methylation, and to identify the main drivers of epigenetic diversity across 22 vineyards planted with the cultivar Shiraz in six distinctive wine areas of a major wine zone, The Barossa, South Australia. The observed epigenetic profiles showed a high level of differentiation that grouped vineyards by their area of provenance despite the low genetic differentiation between vineyards and sub-regions. Furthermore, pairwise epigenetic distances between vineyards with similar management systems showed a significant correlation with geographic distance. Finally, methylation sensitive Genotyping By Sequencing identified 3,598 differentially methylated genes that were assigned to 1,144 unique GO terms of which 8.6% were associated with response to environmental stimulus. Taken together, our results indicate that the intensity and directionality of DNA methylation differentiation between vineyards and wine sub-regions within The Barossa are driven by management and local growing conditions. Finally, we discuss how epigenetic variability can be used as a tool to understand and potentially modulate terroir in grapevine.

## Introduction

The ability of plants to produce alternative phenotypes in response to changing environments is known as phenotypic plasticity (Pigliucci, 2005). Genotypes with this characteristic are able to produce a variety of phenotypes including improved growth and reproduction (Lacaze et al., 2009). Grapevine (*Vitis vinifera* L.) is a highly-plastic crop that exhibits large differences in fruit quality from a given variety depending upon the environmental conditions of the vineyard of origin (Dal Santo et al., 2016). Fruit traits that affect wine quality are thought to be largely driven by the interaction of a grapevine’s genetic characteristics with environmental factors (i.e. climate, soil and topography) and vineyard management (Robinson et al., 2012). All these variables, in conjunction with the wine making process, give a wine its distinctive character. The impact of the environment on grape quality and subsequent wine excellence has given rise to the concept of ‘terroir’, a French term referring to *terre*, “land” (Fanet and Brutton, 2004).

Terroir is defined as the interaction between the physical and biological environment and applied viticultural and oenological practices that lead to unique characteristics in a final wine (Seguin, 1986). Extensive studies have been published on terroir, but generally, these focus on a single parameter such as climatic factors, soil structure or soil microbiology (Harrison, 2000; Tonietto and Carbonneau, 2004;). However, studying only one environmental parameter does not provide an entire understanding of how wine quality is influenced by terroir (Van Leeuwen et al., 2004). A significant amount of work has also been published on the genetic basis of fruit quality in grapevines (eg. Doligez et al, 2002). Despite these insights, further research is required on the molecular changes that are involved in the vine interaction with its environment.

One of the molecular changes worth investigating relates to environmentally induced epigenetic modifications. In fact, phenotypic plasticity has been previously associated to epigenetic variation (Vogt, 2015). Interestingly, analysis of epigenetic diversity has been shown to be more effective in discriminating inter-clonal variability in grapevine than the use of purely genetic molecular markers such as simple sequence repeats (SSRs) or amplified fragment length polymorphisms (AFLPs) (Imazio et al., 2002; Schellenbaum et al., 2008; Ocaña et al., 2013).

Epigenetic mechanisms refer to molecular changes that affect gene expression without changing the organism DNA sequence (Jaenisch and Bird, 2003; Haig, 2004). Epigenetic mechanisms act as an interface between the environment and the genotype by regulating gene expression in response to development and environmental cues and, ultimately affect the plant’s phenotype (Tricker et al., 2012; Kumar et al., 2016). Epigenetic priming is an adaptive strategy by which plants use their memory of the environment to modify their phenotypes to adapt to subsequent conditions (Kelly et al., 2012; Tricker et al., 2013a; 2013b, Vogt, 2015). It is now also widely accepted that epigenetic mechanisms have been the source of useful variability during crop varietal selection (Amoah et al., 2012; Bloomfield et al., 2014; Rodríguez López and Wilkinson, 2015). Of the known epigenetic mechanisms, cytosine methylation (5mC) is arguably the best understood (Goldberg et al., 2007). In plants, 5mC occurs at different cytosine contexts (CpG, CpHpG or CpHpH) (H = A, T or C) (Richards, 1997) and it is induced, maintained or removed by different classes of methyltransferase in conjunction with environmental and developmental cues (Jaenisch and Bird, 2003). It is commonly accepted that DNA methylation constitutes an adaptation strategy to the environment (YunLei et al., 2009), and that changes in DNA methylation can produce altered phenotypes (Zhang et al., 2006; Herrera and Bazaga, 2011; Iqbal et al., 2011). To this extent, there have been extensive studies establishing a link between DNA methylation in plants and environmental conditions (Fonseca Lira-Medeiros et al., 2010; Herrera and Bazaga, 2010; Alonso et al., 2016).

The contribution of DNA methylation to plant performance has been extensively studied in model organisms and some annual crops (Rodríguez López and Wilkinson, 2015). However, we are only beginning to understand how long-living plants, such as grapevines, use epigenetic mechanisms to adapt to changing environments (Fortes and Gallusci, 2017). Effects of environmental conditions on non-annual crops performance can be very difficult to evaluate since many environmental factors interact over the life of the plant to ultimately contribute towards the plant’s phenotype (Fortes and Gallusci, 2017). Although epigenetic mechanisms have been shown to act as a system that allows information storage in organisms across all kingdoms (Levenson and Sweatt, 2005), very few studies have focussed on DNA methylation changes in grapevine. The few known studies in this field used Methylation Sensitive Amplified Polymorphisms (MSAPs) (Reyna-López et al., 1997) for the detection of *in vitro* culture induced epigenetic somaclonal variability (Baránek et al., 2015), and the identification of commercial clones (Imazio et al., 2002; Schellenbaum et al., 2008; Ocaña et al., 2013). However, these studies did not investigate the molecular drivers of terroir.

In this study, we hypothesize that DNA methylation can play a role in defining terroir. To test this hypothesis we investigated the effect of environmental and management conditions on DNA methylation variation in grapevine cultivar Shiraz across 22 vineyards representative of The Barossa wine zone (Australia) (Robinson and Sandercock, 2014) using MSAPs. Finally, we used methylation sensitive Genotyping By Sequencing to characterize the genomic context of the observed regional genetic and epigenetic variability.

## Material and Methods

### Vineyard selection and plant material

Vines from 22 commercial vineyards located in the iconic Barossa wine zone (The Barossa hereafter) (Australia) were included in this study. Vineyards were selected to be representative of the two Barossa Regions as described by the Barossa Grounds Project (Robinson and Sandercock, 2014) (i.e. Eden Valley (three vineyards) and Barossa Valley (19 vineyards which included vineyards in the five distinctive sub-regions within the Barossa Valley Region: Northern Grounds (four vineyards), Central Grounds (four vineyards), Eastern Edge (four vineyards), Western Ridge (four vineyards), Southern Grounds (three vineyards)) (Table S1). To simplify the nomenclature, the Eden Valley region, Northern, Central, Southern Grounds, Eastern Edge and Western Ridge will be defined as sub-regions hereafter. All vineyards were planted with own-rooted vines of the cv Shiraz. Ten vineyards were planted with clone SA 1654 (Whiting, 2003), one with clone BVRC30 (Whiting, 2003), one with clone PT15 Griffith (Farquhar, 2005) and 10 of ‘unknown’ clonal status (Table S1).

A total of 198 plants (nine plants per vineyard) were selected to capture the diversity from each vineyard. Leaf samples (first fully expanded leaf at bud burst, E-L 7) (Coombe, 1995) were collected from three nodes per plant and pooled into a single sample per plant. All samples were taken before dawn (between 10:00 pm and sunrise) to minimize variability associated with differences in plant water status (Williams and Araujo, 2002). Samples were immediately snap-frozen in liquid nitrogen in the vineyard and stored at -80°C until DNA extraction.

### DNA Isolation

Genomic DNA (gDNA) extractions from all 198 samples were performed using the three pooled leaves per plant powdered using an automatic mill grinder (Genogrinder). The obtained frozen powder was used for DNA extraction using the Oktopure automated DNA extraction platform (LGC Genomics GmbH) according to the manufacturer’s instructions. Isolated DNA was quantified using the Nanodrop 2000 spectrophotometer (Thermo Fisher Scientific, Wilmington, USA). DNA final concentrations were normalised to 20 ng/μl using nanopure water (Eppendorf, Germany).

### Analysis of genetic/epigenetic variability using MSAP

MSAP analysis was performed as described by Rodríguez López *et al.*, (2012). In brief, genomic DNA from 88 plants (four plants per vineyard) was digested with a combination of the restriction enzymes *Eco*RI and one of two DNA methylation sensitive isoschizomers (*Hpa*II or *Msp*I). Double stranded DNA adapters (See Table S2 for the sequence of all oligonucleotides used) containing co-adhesive ends complementary to those generated by EcoRI and *Hpa*II/*Msp*I were ligated to the digested gDNA and then used as a template for the first of two consecutive selective PCR amplifications in which the primers were complementary to the adaptors but possessed unique 3’ overhangs. The second selective PCR amplification used primers containing 3’ overhangs previously tested on grapevine (Baránek et al., 2015). *Hpa*II/*Msp*I selective primer was 5’ end-labelled using a 6-FAM reporter molecule for fragment detection using capillary electrophoresis on a ABI PRISM 3130 (Applied Biosystems, Foster City, CA) housed at the Australian Genome Research Facility Ltd, Adelaide, South Australia.

Generated electropherograms were visualized using GeneMapper Software v4 (Applied Biosystems, Foster City, CA). A binary matrix containing presence (1) absence (0) epilocus information was generated for each enzyme combination (i.e. *Eco*RI/*Hpa*II and *Eco*RI/*Msp*I). MSAP fragment selection was limited to allelic sizes between 95 and 500 bp to reduce the potential impact of size co-migration during capillary electrophoresis (Caballero et al., 2008). Different levels of hierarchy were used to group the samples. Samples were first grouped according to vineyard of origin. Then, samples were divided into their sub-regions of origin. Finally, samples were further separated into groups according to clones and the vineyard management systems (i.e. pruning system used in their vineyard of origin) (Table S1).

*Hpa*II and *Msp*I binary matrices were then used to compute Shannon’s Diversity Index implemented using *msap* R package (v. 1.1.8) (Perez-Figueroa, 2013) and Principal Coordinate Analysis (PCoA) was estimated in all regions to determine and visualize the contribution to the observed molecular variability within regions of non-methylated polymorphic loci (NML) and of methylation sensitive polymorphic loci (MSL) (genetic and epigenetic variability respectively) (Smouse et al., 2015).

GenAlex v 6.5 software (Peakall and Smouse, 2006) was used for Principal Coordinate Analysis PCoA in order to visualise the molecular differentiation between Barossa sub-regions detected using MSAP profiles generated after the restriction of gDNA with *Hpa*II or *Msp*I. We then used analysis of molecular variance (AMOVA) to determine the structure of the observed variability using PCoA. Molecular differences between vineyards and regions was inferred as pairwise PhiPT distances (Michalakis and Excoffier, 1996).

Mantel test analysis (Hutchison and Templeton, 1999) was used to estimate the correlation between the calculated pairwise molecular distances with 1. the geographic distance (GeoD) (i.e. Log(1+GeoD (km)) and 2. differences in environmental variables among vineyards (i.e, vineyard altitude, regional average annual rainfall, regional growing season rainfall, regional mean January temperature, regional growing season temperature and growing degree days). Mantel test was implemented in Genalex v 6.5 as described by Rois et al., (2013) and significance was assigned by random permutations tests (based on 9,999 replicates).

### Characterization of genetic/epigenetic variability using msGBS

msGBS was performed as described by Kitimu et al (2015). In brief, 200ng of genomic DNA from 9 samples from Northern, Central and Southern Grounds (vineyards 1-4, 5-8 and 13-15 respectively) were digested using 8U of HF-*Eco*RI and 8 U of *Msp*I (New England BioLabs Inc., Ipswich, MA, USA) in a volume of 20 μl containing 2 μl of NEB Smartcut buffer at 37◸for 2 h followed by enzyme inactivation at 65◸for 10 min. Sequencing adapters were ligated by adding 0.1 pmol of the *Msp*I adapters (uniquely barcoded for each of the 198 samples) and 15 pmol of the common *Eco*RI Y adapter (See Table S2 for the sequence of all oligonucleotides used), 200 U of T4 Ligase and T4 Ligase buffer (New England BioLabs Inc., Ipswich, MA, USA) in a total volume of 40 μl at 24◸for 2 h followed by an enzyme inactivation step at 65◸for 10 min. Excess adapters were removed from ligation products using Agencourt AMPure XP beads (Beckman Coulter, Australia) at the ratio of 0.85 and following manufacturer’s instructions. Single sample msGBS libraries were then quantified using Qbit 3 (Thermofisher). A single library was generated by pooling 25 ng of DNA from each sample. Library was then amplified in 8 separate PCR reactions (25 μl each) containing 10 μl of library DNA, 5 μl of 5x Q5 high fidelity buffer, 0.25 μl polymerase Q5 high fidelity, 1 μl of each forward and reverse common primers at 10 μM, 0.5 μl of 10 μM dNTP and 7.25 μl of pure sterile water. PCR amplification was performed in a BioRad T100 thermocycler consisting of DNA denaturation at 98°C (30 s) and 10 cycles of 98°C (30 s), 62°C (20 s) and 72°C (30 s), followed by 72°C for 5 minutes. PCR products were then re-pooled and DNA fragments ranging between 200 and 350 bp in size were captured using the AMPure XP beads following manufacturer’s instructions. Libraries were sequenced using an Illumina NextSeq High Output 75bp pair-end run (Illumina Inc., San Diego, CA, USA) at the Australian Genome Research Facility (AGRF, Adelaide, Australia).

### msGBS Data analysis

Analysis of genetic diversity between regions was performed by single nucleotide polymorphism (SNP) calling using TASSEL 3 (Bradbury et al, 2007) on msGBS sequencing results. Only SNPs present in at least 80% of the samples were considered for analysis. Principal component analysis (PCA) was implemented on TASSEL 3 using the selected SNPs. To identify any possible geographical genetic structure, the optimal number of genetic clusters present in the three regions were computed using Bayesian Information Criterion (BIC) as effected by Discriminant Analysis of Principal Components (DAPC) using adegenet 2.0.0 (http://adegenet.r-forge.r-project.org/).

Identification of significant differentially methylated markers (DMMs) between regions was then computed using the package *msgbsR* (https://github.com/BenjaminAdelaide/msgbsR), accessed on 26/08/2016). In brief, raw sequencing data was first demultiplexed using GBSX (Herten et al, 2015) and filtered to remove any reads that did not match the barcode sequence used for library construction. Following demultiplexing, paired-end reads were merged using bbmerge in bbtools package (Bushnell, 2016). Merged reads were next aligned to the 12X grapevine reference genome (http://plants.ensembl.org/Vitis_vinifera/). Alignment BAM files where then used to generate a read count matrix with marker sequence tags, and used as source data to perform subsequent analyses using *msgbsR* in the R environment (R Core Team, 2015). Finally, the presence of differential methylation between regions was inferred from the difference in the number of read counts from all sequenced *Msp*I containing loci that had at least 1 count per million (CPM) reads and present in at least 15 samples per region. Significance threshold was set at Bonferroni adjusted P-value (or false discovery rate, FDR) < 0.01 for difference in read counts per million. The *log*FC (logarithm 2 of fold change) was computed to evaluate the intensity and direction of the region specific DNA methylation polymorphism.

To determine how the observed changes in DNA methylation between sub-regions were associated to protein coding genes, the distribution of DMMs was assessed around such genomic features, as defined in Ensembl database (http://plants.ensembl.org/biomart/martview/), by tallying the number of DMMs between the transcription start site (TSS) and the transcription end site (TES) and within five 1 Kb windows before the TSS and after TES of all *V. Vinifera* genes, using *bedtools* (Quinlan and Hall, 2010).

Genes within 5 Kb of a DMM were referred to as differentially methylated genes (DMGs). DMGs in each pairwise regional comparison were grouped into those showing hypermethylation or hypomethylation, and were next used separately for GO terms enrichment, using the R packages: *GO.db* (Carlson, 2016) and *annotate* (Gentlemen, 2016). Significant GO terms were selected based on Bonferroni adjusted P-values at significance threshold of 0.05. Finally, GO terms containing DMGs in all three pairwise comparisons were visualized using Revigo (Supek et al, 2011).

## Results

Analysis of MSAP profiles obtained from DNA extractions of the first fully expanded leaf of 88 individual vines selected from 22 commercial vineyards within the six Barossa sub-regions (Figure 1; Table S1) yielded 215 fragments comprising 189 from *Msp*I and 211 from *Hpa*II, of which 80% and 84% respectively, were polymorphic (i.e. not present in all the analysed samples/replicates when restricted with one of the isoschizomers).

**Figure 1:**
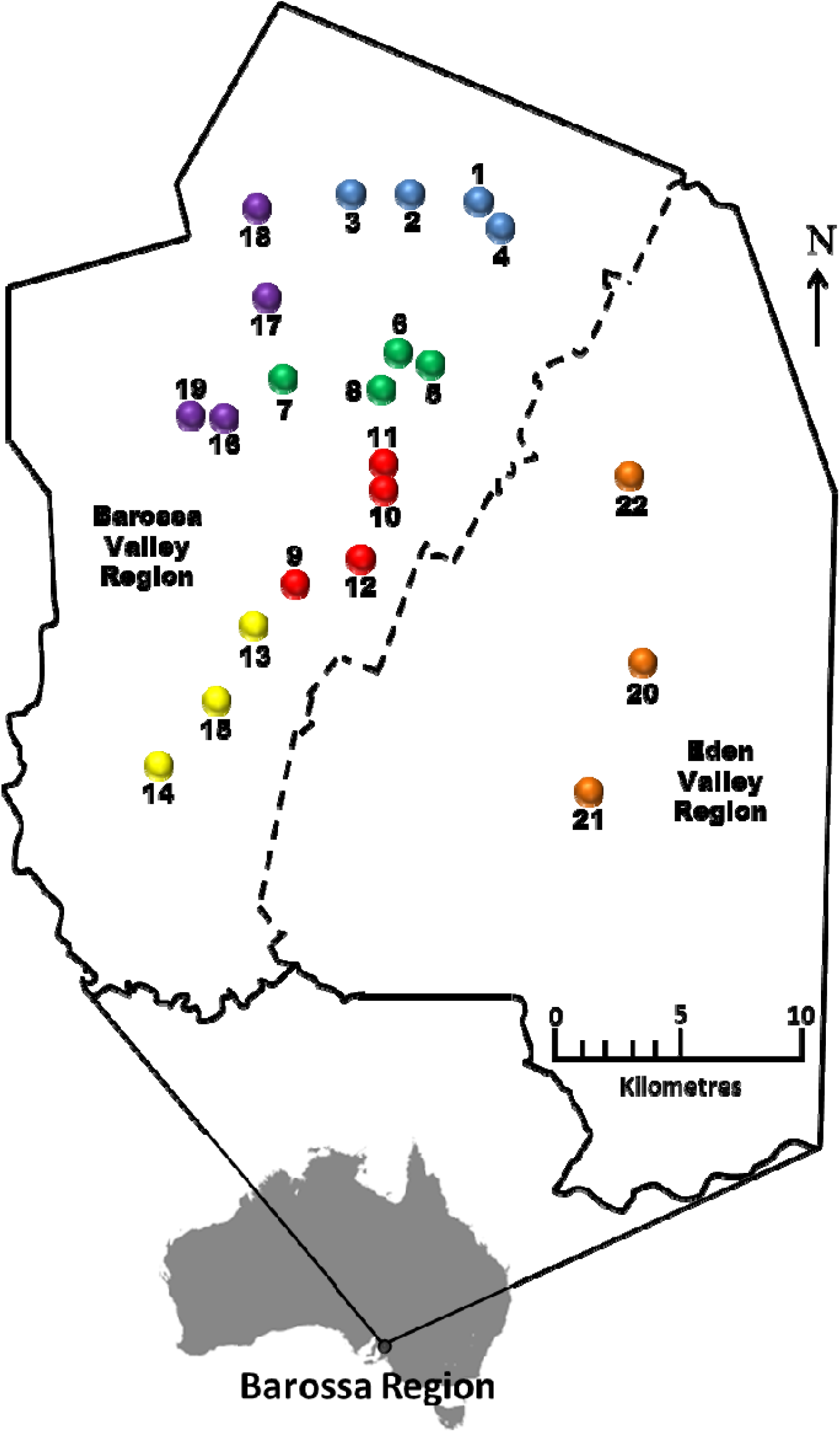
Selected Barossa Region vineyard sites. Northern Grounds: Blue, Southern Grounds: Yellow, Central Grounds: Green, Eastern Edge: Red, Western Ridge: Purple, Eden Valley: Orange. Arrow indicates geographic north.

### Analysis of genome/methylome differences within wine sub-regions in The Barossa

PCoA of the MSAP profiles generated from non-methylated polymorphic loci (NML) (genetic variability) and by methylation sensitive polymorphic loci (MSL) (epigenetic variability) (Pérez-Figueroa, 2013) revealed a higher separation between vineyards when using epigenetic information than when using genetic (Figure S1). The capacity of both types of variability to differentiate between vineyards was more evident on vineyards in the Southern Grounds (Figure 2C-D). Both PCoA analysis and Shannon’s diversity index showed significantly higher epigenetic than genetic diversity for all sub-regions (Figure S2, Table 1). Among sub-regions, Southern Grounds had the highest epigenetic diversity (0.581 ± 0.124) and Western Ridge the lowest (0.536 ± 0.143). Genetic diversity showed the highest value in the Southern Grounds (0.374 ± 0.143) and the lowest in the Northern Grounds (0.240 ± 0.030).

**Figure 2.**
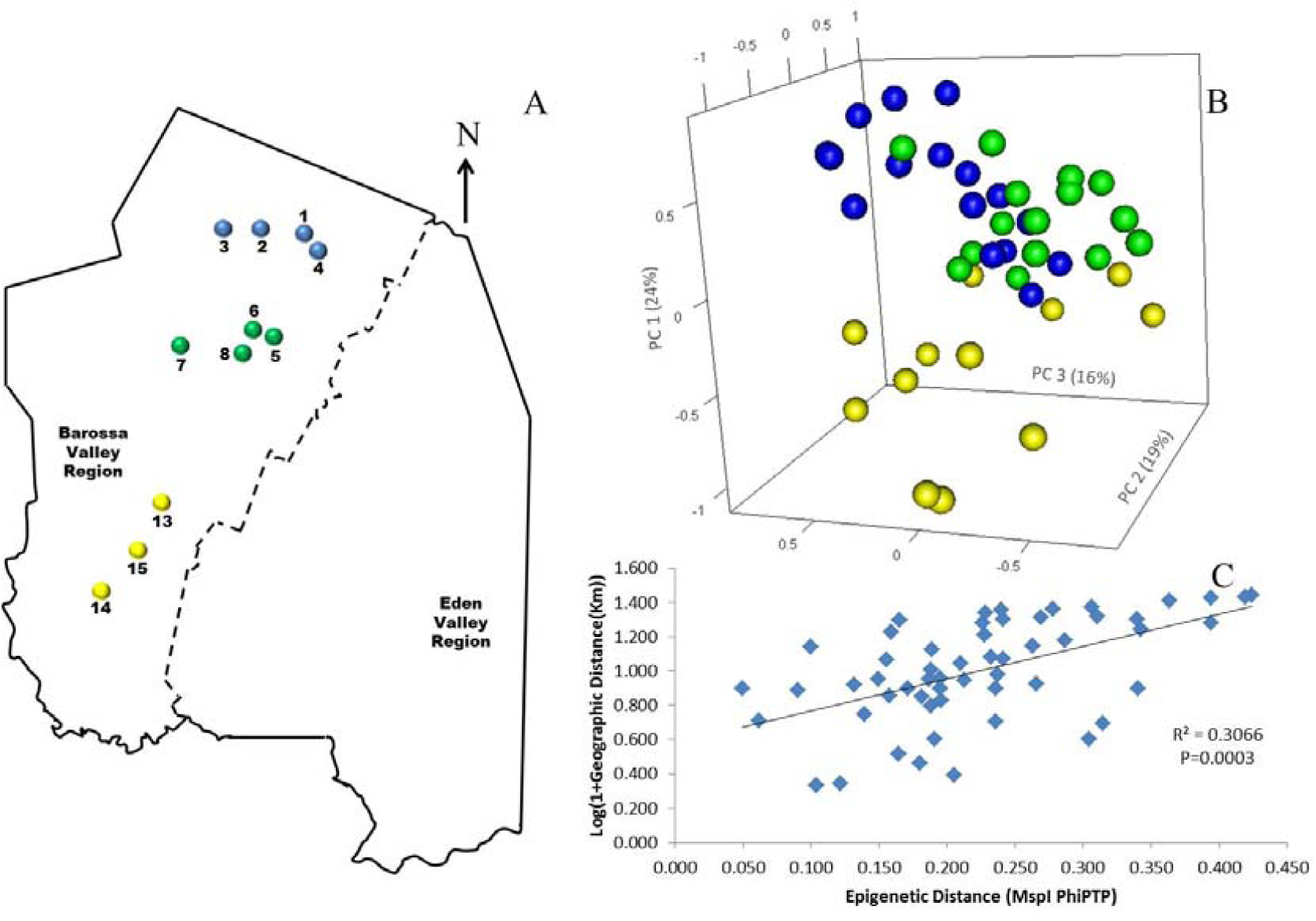
Analysis of the correlation between epigenetic differentiation and geographic distance of vineyards planted along the Barossa Valley North-South axis: **(A)** Location of the Barossa Valley vineyards from the three sub-regions distributed along the Barossa Valley North-South axis; Northern Grounds (blue), Central Grounds (Green) and Southern Grounds (Yellow). Arrow indicates the direction of geographic North. **(B)** Principal Coordinate Analysis (PCoA) representing epigenetic differences between leaf samples collected from 4 plants/vineyard. Percentage of the variability capture by each Principal Coordinate (PC) is shown in parenthesis. **(C)** Correlation between pairwise epigenetic distance (*Msp*I PhiPT) and geographical distance (Log(1+GeoD (km)) between vineyards. Shown equations are the correlation’s R^2^ and the Mantel test significance (P value was estimated over 9,999 random permutations tests). PCoA and PhiPT for Mantel test were based on presence/absence of 215 loci obtained from MSAP profiles generated using *Msp*I.

**Table 1.**
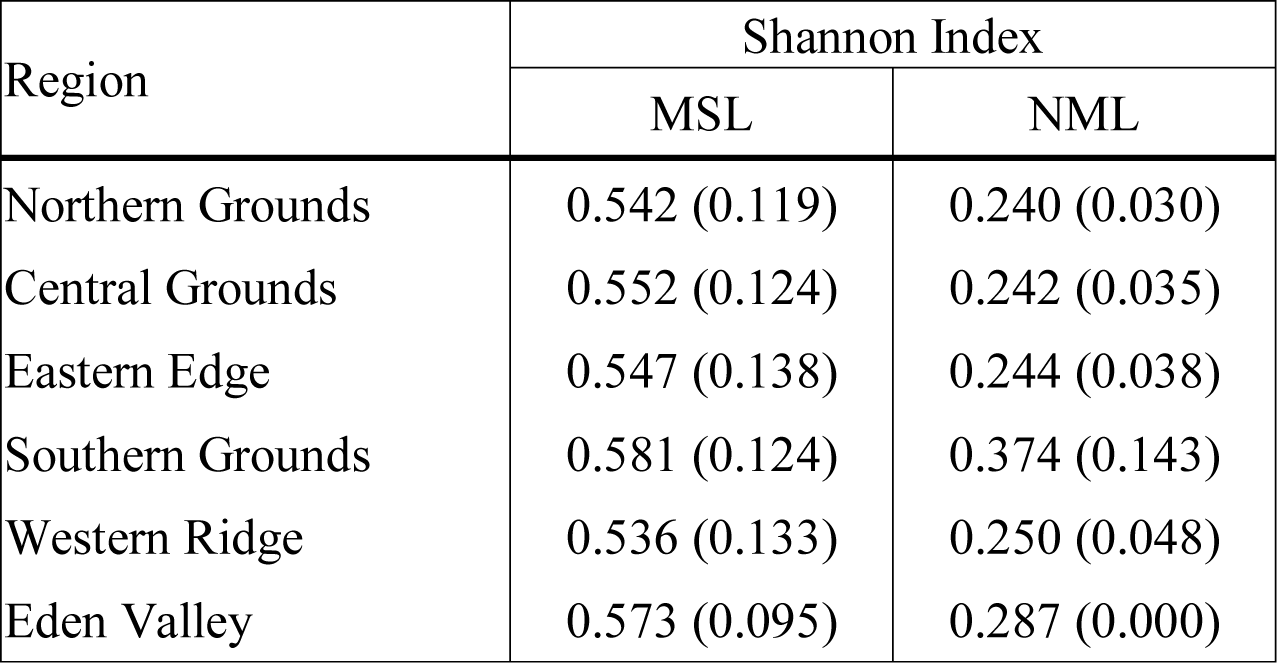
Analysis of genetic (NML) and epigenetic (MSL) diversity within the six Barossa Valley wine growing regions: Shannon diversity indices are reported as mean (± Standard Deviation). Wilcoxon rank test provides statistical support for all Shannon diversity indices (P< 0.0001).

### Analysis of genome/methylome differences between wine sub-regions in The Barossa

We used analysis of the molecular variance (AMOVA) (Table 2) to obtain an overview of the molecular variability between all the studied sub-regions. Overall, MSAP profiles generated using restriction enzyme *Msp*I achieved better separation between sub-regions than those generated using *Hpa*II. Of all 30 calculated molecular pairwise distances between sub-regions (PhiPTs), 25 were significant (P<0.05) (Table 2). Calculated PhiPT values ranged from 0.115 (PhiPT of Northern Grounds vs Southern Grounds calculated using *Msp*I) and 0.012 (PhiPT of Central Grounds vs Eastern Edge calculated using *Hpa*II).

**Table 2:** Molecular distances (PhiPT) between Barossa Valley wine producing sub-regions. PhiPT values were calculated using MSAP profiles generated from 88 grapevine plants (4 individual plants per vineyard) using restriction enzyme combinations *Msp*I/*EcoR*I (above diagonal) and *Hpa*II/*EcoR*I (below diagonal). P-values (shown in parenthesis) were calculated based on 9,999 permutations. Pairwise regional comparisons showing not significantly different PhiPT values are highlighted in bold. A total of 22 vineyards were included in the analysis: Northern Grounds (4), Central Grounds (4), Southern Grounds (3), 4 vineyards in Eastern Edge (4), 4 vineyards in Western Ridge (4) and Eden valley (3).

|  | North | South | Central | East | West | Eden |
| --- | --- | --- | --- | --- | --- | --- |
| North | – | 0.115(1e-04) | 0.043(8e-04) | 0.062(2e-04) | 0.082(1e-04) | 0.069(0.001) |
| South | 0.028(0.0059) | – | 0.064(1e-04) | 0.042(0.001) | 0.027(0.024) | **0.024(0.073)** |
| Central | **0.012(0.1085)** | 0.025(0.0079) | – | 0.043(1e-04) | 0.060(1e-04) | 0.067(2e-04) |
| East | 0.025(0.0043) | **0.012(0.0712)** | 0.015(0.0474) | – | 0.029(0.004) | 0.038(0.0011) |
| West | 0.039(2e-04) | 0.018(0.0426) | 0.033(0.0001) | **0.013(0.0651)** | – | 0.024(0.024) |
| Eden | 0.056(0.0001) | 0.043(4e-04) | **0.015(0.0601)** | 0.031(0.0023) | 0.031(0.0016) | – |

AMOVA on MSAP profiles indicates that the majority of the observed variability is explained by differences within vineyards (81% using profiles generated with *Msp*I and 91% with *Hpa*II). A significant proportion of the total variability detected was associated to differences between vineyards (17% with *Msp*I and 8% with *Hpa*II) and 2% and 1% was due to differences between sub-regions (*Msp*I and *Hpa*II respectively).

### Effect of vineyard location on methylome differentiation

To determine if environmental differences between vineyards influenced the observed epigenetic differences we studied the vineyards’ pairwise geographic and molecular distances correlation. Vineyards located on the North-South axis of the Barossa Valley (i.e, vineyards 1, 2, 3 and 4 (Northern Grounds), 5, 6, 7 and 8 (Central Grounds), and 13, 14 and 15 (Southern Grounds)) (Figure 2A) were selected as Northern and Southern Grounds showed the greatest epigenetic differentiation (Table 2). PCoA analysis showed that Central Grounds samples occupied an intermediate Eigen space between Northern and Southern Grounds samples with coordinate 1 (24% of the observed variability) representing the North-South axis (Figure 2B). Moreover, Mantel test showed a significant (P = 0.0003) positive correlation (R^2^=0.3066) between pairwise vineyard epigenetic and geographic distances (Figure 2C). Then, Mantel test analysis was implemented to compare the observed molecular differences against environmental variables. Differences in vineyard altitude showed a small but significant positive correlations (R^2^=0.1615, P=0.013) with PhiPT values between vineyards (Figure S3). We then investigated if clone and vineyard management systems could be contributing to this correlation, by comparing the epigenetic/geographic distances correlation of 10 vineyards planted with clone 1654 (vineyards 1 and 4 (Northern Grounds), 7 (Central Grounds), 9 and 12 (Eastern Ridge), 15 (Southern Grounds) 16, 17, 18 and 19 (Western Ridge) (Figure 3A) and of 6 vineyards planted with the same clone (1654) and trained using the same pruning system (i.e. spur pruning) (vineyards 1 (Northern Grounds), 7 (Central Grounds), 9 (Eastern Ridge), 15 (Southern Grounds) 16 and 19 (Western Ridge) (Figure 4A)). Again, PCoA shows that the main contributor (23-24%) to the detected variability is associated to the distribution of the vineyards on the N-S axis. Mantel test showed a positive correlation for both epigenetic/geographic distance comparisons, however, although both correlations were significant (P< 0.05), the correlation among vineyards pruned using the same system (Figure 4B-C) was higher than that observed when all pruning systems were incorporated in the analysis (Figure 3B-C).

**Figure 3.**
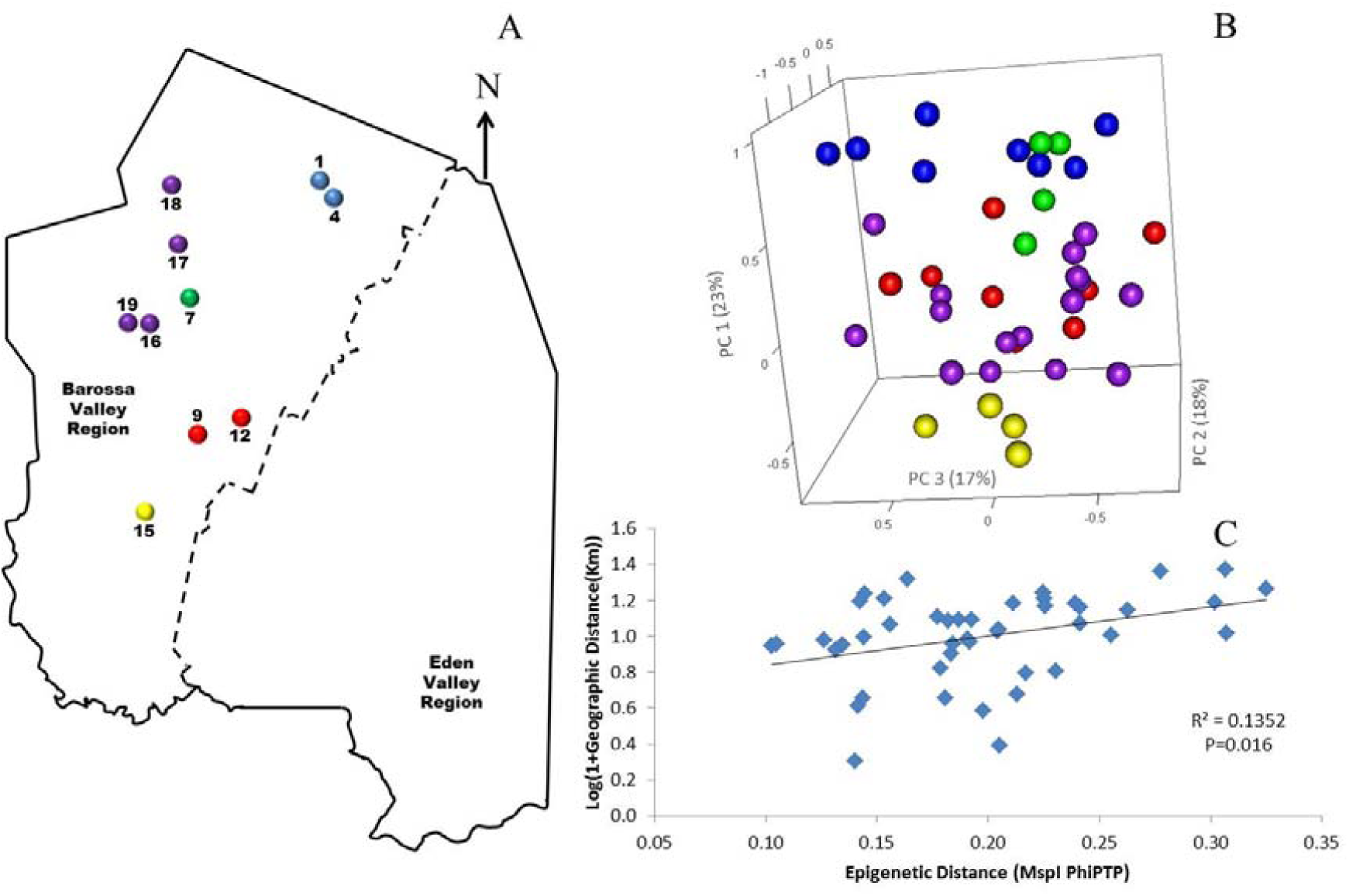
Analysis of the correlation between epigenetic differentiation and geographic distance of vineyards planted with clone 1654 in the Barossa Region: **(A)** Location of the selected Barossa Valley vineyards from the three sub-regions distributed along the Barossa Valley North-South axis Northern Grounds (blue), Central Grounds (Green), Eastern Edge (red), Southern Grounds (Yellow) and Western Ridge (Purple). Arrow indicates the direction of geographic North. **(B)** Principal Coordinate Analysis (PCoA) representing epigenetic differences between leaf samples collected from 4 plants/vineyard. Percentage of the variability captured by each Principal Coordinate (PC) is shown in parenthesis. **(C)** Correlation between pairwise epigenetic distance (*Msp*I PhiPT) and geographical distance (Log(1+GeoD (km)) between vineyards. Shown equations are the correlation’s R^2^ and the Mantel test significance (P value was estimated over 9,999 random permutations tests). PCoA and PhiPT for Mantel test were based on presence/absence of 215 loci obtained from MSAP profiles generated using *Msp*I.

### msGBS analysis of genome/methylome differentiation between Northern, Central and Southern Grounds

TASSEL 3 was then implemented on msGBS data for single nucleotide polymorphism (SNP) calling from 99 samples collected in 11 vineyards in the Northern, Central and Southern Grounds sub-regions. This generated a total of 8,139 SNPs of which 4,893 were present in at least 80% of the sequenced samples. PCA analysis using filtered SNPs showed very low level of genetic structure, with only five plants from vineyard 3 (Northern Grounds) separating from the rest (Figure S4A). However, this clustering was not supported by DAPC (i.e. the optimal clustering solution should correspond to the lowest BIC) which indicated the optimal number of clusters for this data set is 1 (Figure S4B) suggesting a lack of genetic structure in the vineyards/regions analysed.

PC-LDA analysis was then used to visualize differences in DNA methylation detected using msGBS. DNA methylation profiles clustered samples by their sub-region of origin, with Northern and Central Grounds being separated by differential factor (DF1) from Southern Grounds while DF2 separated Northern from Central Grounds (Figure 5). These results were supported by the higher number of DMMs found when comparing samples from Southern to samples from Central or Northern Grounds than when comparing Northern to Central (Table 3).

**Figure 5.**
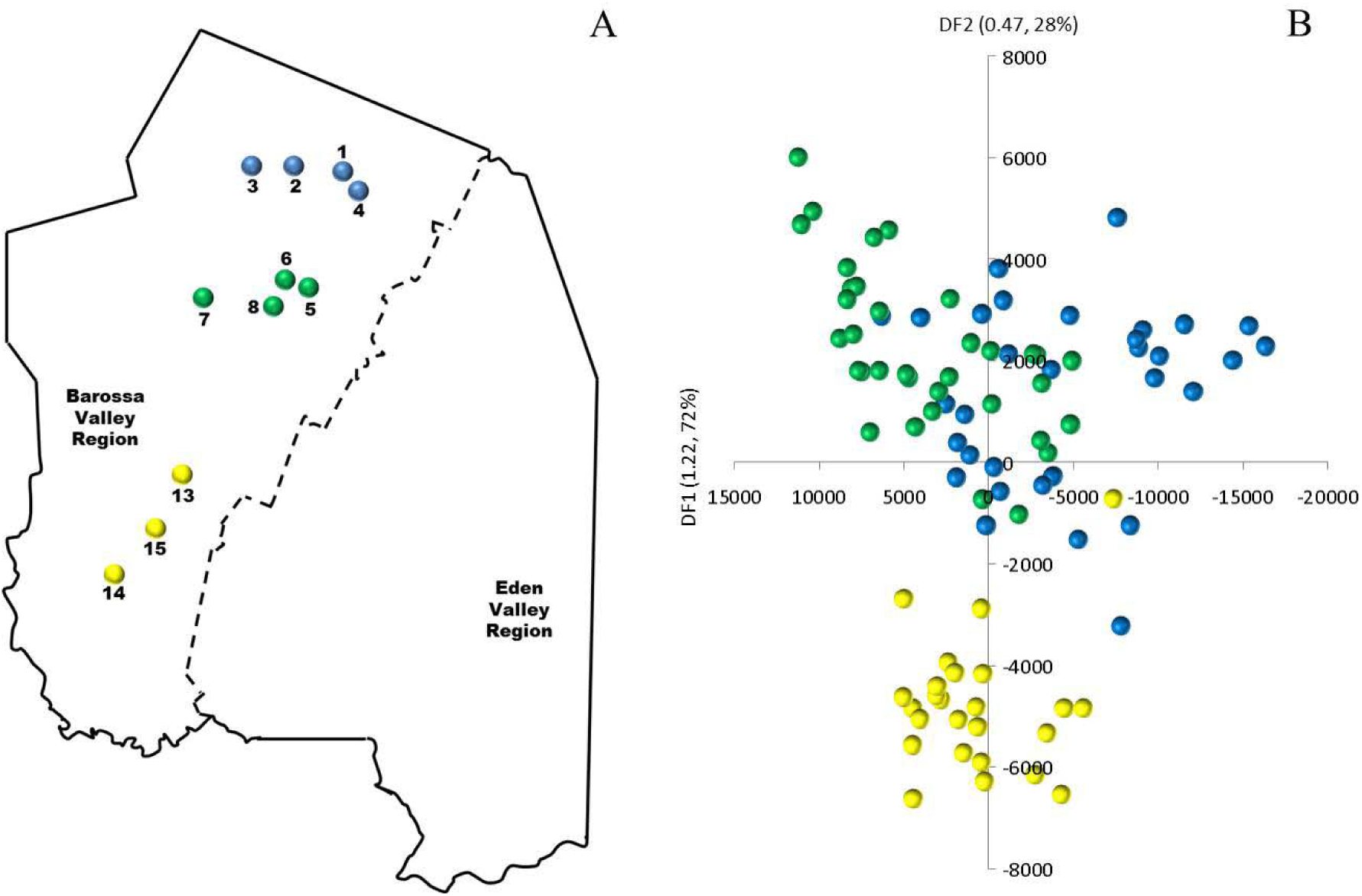
Analysis of the correlation between epigenetic differentiation and geographic distance of vineyards planted along the Barossa Valley North-South axis: **(A)** Location of the Barossa Valley vineyards from the three sub-regions distributed along the Barossa Valley North-South axis; Northern Grounds (blue), Central Grounds (Green) and Southern Grounds (Yellow). Arrow indicatesthe direction of geographic North. **(B)** Principal Components-Linear Discriminant Analysis (PC-LDA) representing epigenetic differences between leaf samples collected from 9 plants/vineyard. Percentage of the variability capture by each differential factor (DF) is shown in parenthesis. PC-LDA were based on read number of loci obtained from msGBS profiles.

**Table 3:** Identification of differentially methylated markers (DMMs), genes (DMGs) and GO terms (DMGOs) between sub-regions in Barossa Shiraz. Cells contain the number of DMMs, DMGs or DMGOs detected in each pairwise comparison. Differential methylation between sub-regions was calculated using msGBS data from 9 Shiraz plants per vineyard (Northern Grounds (NG): four vineyards, Central Grounds (CG): Four vineyards and Southern Grounds (SG); Three vineyards). Directionality of methylation (i.e. Hyper/hypomethylation) indicates an increase or decrease in DNA methylation in the second region compared to the first region in each pairwise comparison.

|  | Differentially methylated markers | Differentially methylated genes |  | Differentially methylated GO terms |  |
| --- | --- | --- | --- | --- | --- |
|  |  | Hypometh Genes | Hypermeth Genes | Hypometh GO terms | Hypermeth GO terms |
| **NG vs CG** | 7465 | 374 | 178 | 327 | 204 |
| **SG vs CG** | 15276 | 691 | 2382 | 520 | 832 |
| **SG vs NG** | 12911 | 522 | 2094 | 500 | 815 |

We next investigated the association of the detected DMMs to annotated protein-coding genes in the grapevine genome by surveying their location and density within and flanking such genomic features. A total of 3,598 genes were deemed differentially methylated (i.e. presented one or more DMMs within 5kb of the TSS or the TSE) or within genes (Table 3). Quantification of such DNA methylation changes showed that, in average, methylation levels are higher in the northern most region in each comparison (i.e. NG > CG > SG) (Figure 6A). The majority of detected DMMs associated to a gene were present in the body of the gene and the number of DMMs decreased symmetrically with distance from the TSS and the TES (Figure 6B and Tables S3, S4 & S5). Finally, as observed with all DMMs, the comparison between Northern and Central Grounds samples showed the lowest number of DMGs (Table 3, Figure 6C and Supplementary Tables S3, S4 & S5).

**Figure 6.**
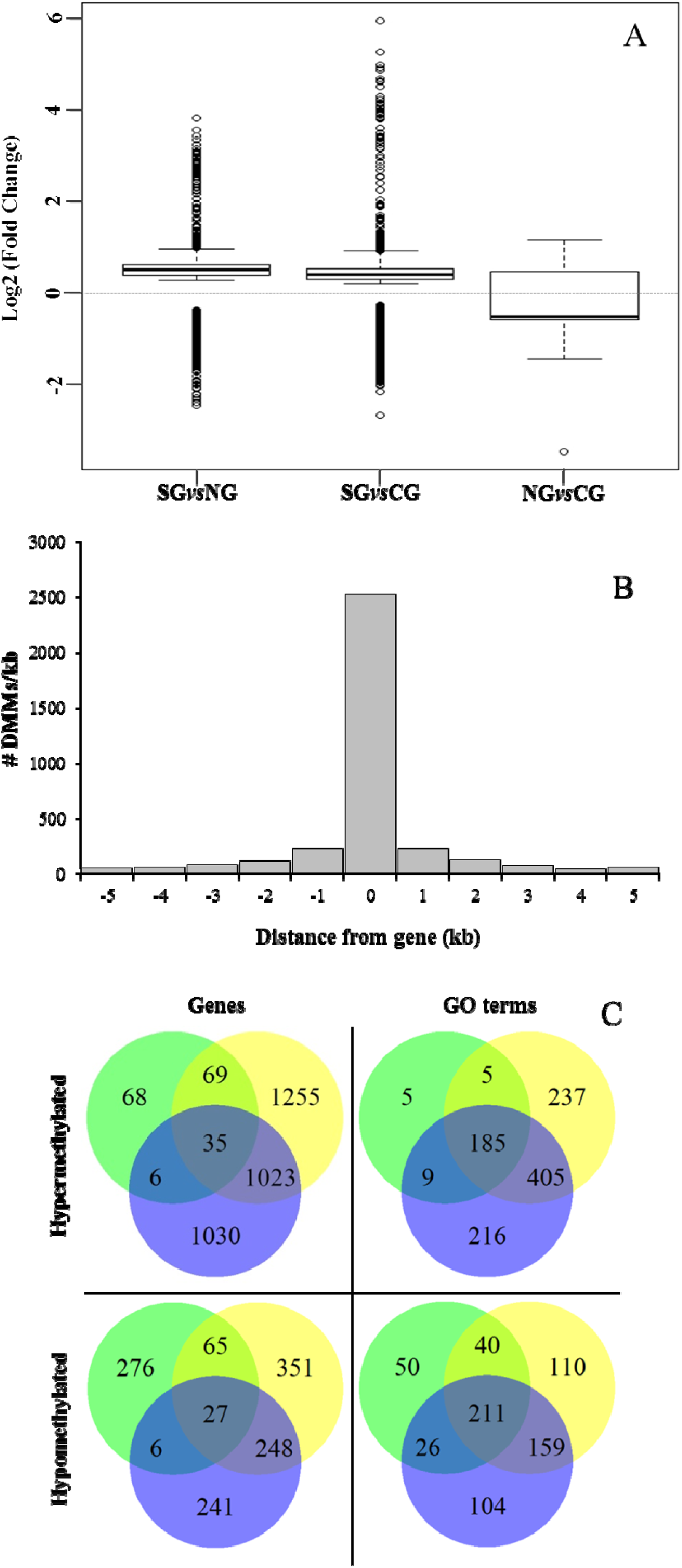
Analysis of differentially methylated genes (DMGs) and GO terms (DMGOs) among three wine sub-regions in Barossa Shiraz. Genes were considered differentially methylated if located within 5kb of at least one differentially methylated marker (DMM) (FDR < 0.01). DMMs were generated using msGBS on 9 plants per vineyard (Northern Grounds: four vineyards, Central Grounds: four vineyards and Southern Grounds; three vineyards). **(A)** Directionality of methylation differences between regions. Boxplots show the distribution of the intensity of changes in DNA methylation level between regions, represented here as the fold-change (2 power *log*2FC) in read counts for a given msGBS markers between two regions. Median shows the direction of the methylation flux at a whole genome level in each region comparison (i.e. positive medians indicate a global increase in DNA methylation (hypermethylation) while negative medians indicate a global decrease in DNA methylation (hypomethylation) in the second region in the comparison (e.g. Northern Grounds is hypermethylated compared to Southern Grounds). **(B)** Distribution of 3598 region specific DMMs around genes. Columns -5 to -1 and 1 to 5 represents the number of DMMs per Kb around *V. vinifera* genes. Column 0 indicates the number of DMMs within the coding sequence (i.e. between the transcription start and end sites) of *V. vinifera* genes. **(C)** Shared DMGs and differentially methylated Gene Ontology Terms (DMGOs) between regional comparisons. Venn diagrams show the number of unique and shared DMGs and DMGOs between each regional pairwise comparison (i.e. Blue: hyper/hypomethylated genes and GOs in Northern Grounds compared to Southern Grounds; Yellow: in Central Grounds compared to Southern Grounds; and Green: in Central Grounds compared to Northern Grounds).

To gain further insight into the functional implications of the DNA methylation differences detected between sub-regions, we used *GO.db* (Carlson, 2016) and *annotate* (Gentleman, 2016) to assign 1144 unique GO terms to the observed DMGs (Adjusted P value <0.05). As observed with DMMs and DMGs the comparison between Northern and Central Grounds samples showed the lowest number of GO terms containing differentially methylated genes (DMGOs) (Table 3, Figure 6C and Tables S3, S4 & S5). REViGO semantic analysis of GO terms shared by all 3 pairwise regional comparisons (Figure 7) showed an increase of gene enrichment (i.e. a decrease in adjusted P values) with geographic distance (e.g. see Figure 7 for comparisons between Northern Grounds and Southern Grounds (A-B) and Central Grounds and Northern Grounds (C-D). 311 DMGs (8.6% of the total) were allocated in GO terms associated to response to environmental stimulus (161 and 150 abiotic and biotic challenges respectively) (Figure 7, Tables S6 & S7), which included GO terms in the semantic space of plant response to light, temperature, osmotic/salt stress and defence to biotic stimulus.

**Figure 7:**
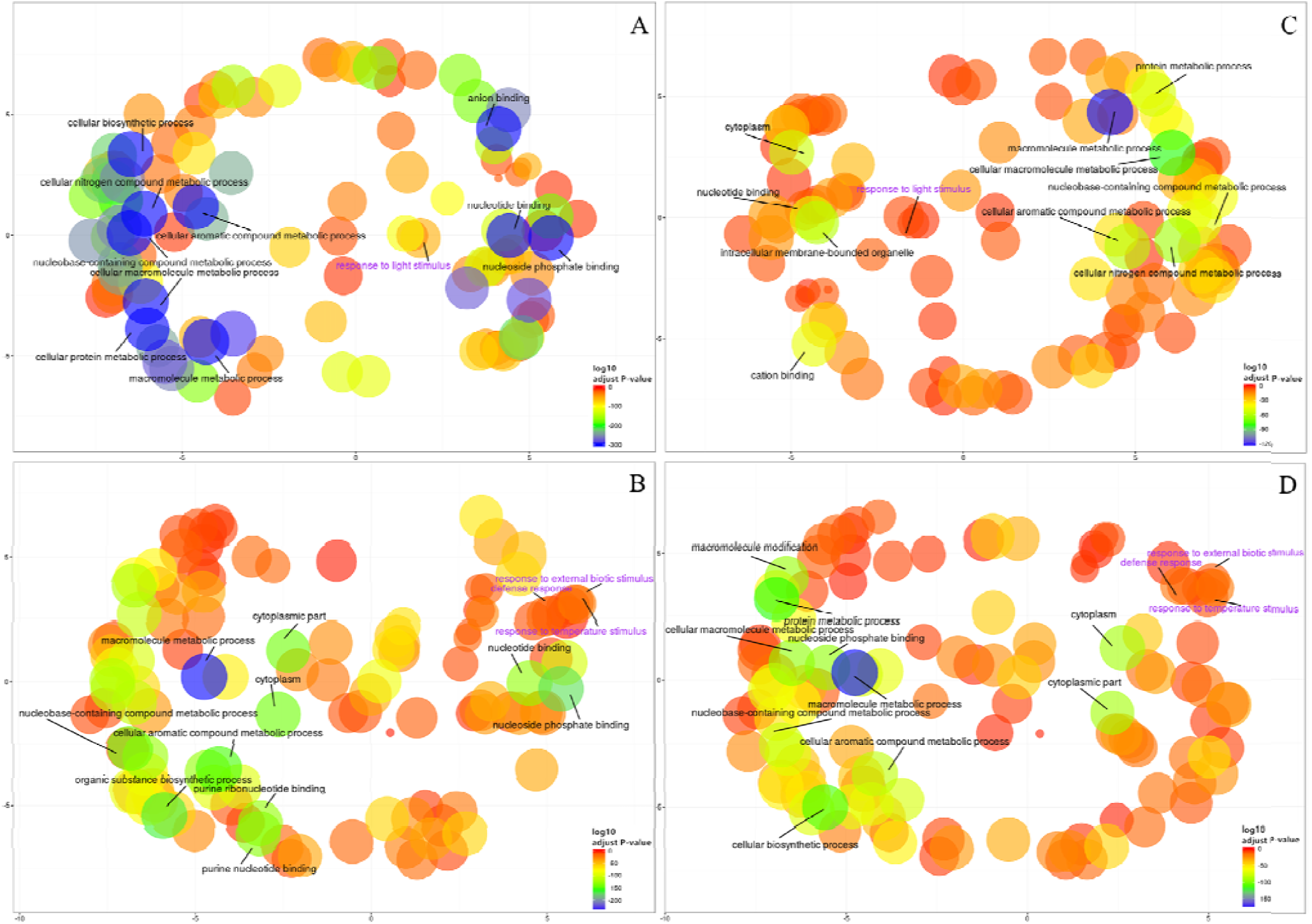
REViGO semantic analysis of differentially methylated GO terms shared by all three regional pairwise comparisons. Functional enrichment of GO-terms was carried out for the genes deemed differentially methylated (DMGs) hypermethylated (185) (A-C) or hypomethylated (211) (B-D) in Northern Grounds compared to Southern Grounds (A-B) and Central Grounds compared to Northern Grounds (C-D) using GO.*db* and *annotate* and summarized using REViGO. Bubble color indicates the p-value for the false discovery rates (the first 10 terms are labelled with legends in black. A detailed list of all GO terms containing DMGs has been supplied as a Tables S6 and S7); circle size indicates the frequency of the GO term in the underlying GO database (bubbles of more general terms are larger).

## Discussion

### Grapevine DNA methylation patterns are region specific

Analysis of *Hpa*II and *Msp*I generated MSAP profiles showed that the methylation profiles of the six different sub-regions were significantly different (P<0.05) in 25 of the 30 possible pairwise comparisons (Table 2). Variability among vineyards and sub-regions was higher in *Msp*I generated profiles (17 and 2%) than in *Hpa*II profiles (8 and 1%), indicating that the detected regional epigenetic differences are, at least partially, sequence context specific (Tricker et al., 2012; Meyer, 2015). Calculated PhiPT values showed low levels of molecular differentiation between sub-regions, even when those differences were statistically significant (Table 2). This could be explained by the high proportion of the total variability associated to differences between individual plants (81-91%) compared to 1-2% associated to differences between sub-regions. Such high levels of molecular differentiation between individuals could be due to the random accumulation of somatic variation with age, which can be genetic or epigenetic in nature. PCA of genetic polymorphisms detected using msGBS results showed a high level of genetic variability between plants (Figure S4A) which is characteristic of long living plants in general (Baali-Cherif and Besnard, 2005) and in grapevine in particular (Torregrosa et al., 2011). However, Discriminant Analysis of Principal Components did not detect any sample clustering associated to their origin (Figure S4B) indicating that genetic diversity is not structured in a geographic manner. Although both genetic and epigenetic somatic variation can be random (Vogt, 2015), different growing conditions will differentially affect the DNA methylation profiles of otherwise genetically identical individuals (Consuegra and Rodríguez López, 2016) as previously shown on clonally propagated *Populus alba* (Guarino et al., 2015). It is, therefore, not surprising to find that epigenetic profiling was a better predictor of sample origin than genetic profiling alone both using MSAP data (Table 2, Figure S1) or msGBS data (Figure 5-7 and Figure S4). This suggests that although genetic differences between regions or vineyards can partly contribute to the observed molecular differentiation between vineyards/sub-regions, epigenetic differences are the major driver of such differentiation.

Samples collected from vineyards in the Southern Grounds presented the highest levels of both genetic and epigenetic diversity (Table 1). These vineyards also presented higher levels of differentiation when inter-vineyard variability was analysed (Figure 2G-H), suggesting a major contributor to the observed molecular variability between vines in the Southern Grounds is linked to the vineyard of origin. Taken collectively, these results suggest that the specific growing conditions from each subregion impose DNA methylation patterns on grapevine plants specific for each region as previously shown both in cultivated (Guarino et al., 2015) and wild plant populations (Fonseca Lira-Medeiros et al., 2010). Not surprisingly, and contrary to what has been shown in natural plant populations (Fonseca Lira-Medeiros et al., 2010; Róis et al., 2013), no clear negative correlation between genetic and epigenetic diversity was observed in the studied vineyards. This is most probably due to the intensive phenotypic selection to which grapevine cultivars have been under since domestication and the relative low levels of environmental disparity to which vines growing in the same vineyard are exposed to.

### Environmental and vineyard management differences are drivers of regional epigenetic differentiation

Principal coordinate and Mantel test analysis showed that the correlation between epigenetic and geographic distance between vineyards on the North-South axis of the Barossa Valley (Figure 2A) was significant (P=0.0003) (Figure 2C) and that the main contributor to the observed epigenetic differences was the position of the studied vineyards along the N-S axis (Figure 2B). This suggests that environmental differences between locations could be contributing to the observed molecular differences between sub-regions or vineyards (Figure 3). Moreover, the correlation (R^2^ = 0.3066) between epigenetic and geographic distance among vineyards planted with clone 1654 on the N-S axis (Figure 2) supports the Shannon diversity analysis that indicate that the different genetic backgrounds used in this study do not greatly affect the epigenetic differences observed between regions (Table 1). Conversely, differences in vineyard altitude appear to be a contributor to the detected epigenetic differentiation between vineyards (Figure S3). Previous work has shown that sun exposure can have significant effects both in berry metabolomic profiles (Son et al., 2009; Tarr et al., 2013) and on the epigenetic profiles of plants growing in different environments (Guarino et al., 2015). Although altitude does not necessarily affect sun exposure, it can have a profound effect on the UV levels experienced by plants (approximately 1% increase every 70 m gain in altitude). Our results indicate that, although DNA methylation in and around genes changes in both directions (hyper-and hypo-methylation), on average, it increases with altitude (i.e. NG > CG > SG (vineyard average altitude 301, 277 and 236 m respectively) (Figure 6A). Functional analysis of the DMGs between sub-regions generated GO terms associated to plant response to light stimulus (Table S7). More importantly, the number of genes associated to such GO terms was higher in comparison between regions with bigger differences in altitude (74 and 46 genes in comparison SG *vs* NG (65 m difference) and SG *vs* CG (41 m) respectively) than in the pairwise comparison with lower difference in altitude (6 genes NG *vs* CG (24 m)). Although this positive polynomial grade 2 correlation (R^2^=1) was generated using only three data points, it is tempting to speculate that differences in light incidence due to differences in altitude are triggering the observed changes in DNA methylation in response to light stimulus genes. Especially when previous work has shown that, in grapevine leaves, increased UV levels trigger the synthesis of non-flavonoid phenolics such as resveratrol (Sbaghi et al., 1995; Teixeira et al., 2013). Interestingly, DNA methylation has been previously linked to the regulation of the gene VaSTS10, which controls the synthesis of resveratrol in *Vitis amurensis* (Tyunin et al., 2013; Kiselev et al., 2013).

To our knowledge the effect of pruning has not yet been studied at an epigenetic level. However Herrera and Bazaga (2011) showed that long term defoliation by herbivory does have an effect on the DNA methylation patterns of predated plants. The correlation between epigenetic and geographic distances observed between vineyards planted with clone 1654 and pruned with the same method (spur pruning) (Figure 4) was reduced when all vineyards planted with clone 1654 were considered irrespectively of the pruning system used (Figure 3). This concerted epigenetic response of plants growing in different environments towards the same human stimulus is consistent with the hypothesis that differences in cultural practices (e.g. pruning method) act together with environmental conditions as major drivers of the epigenetic differences observed between vineyards and sub-regions in this study.

**Figure 4.**
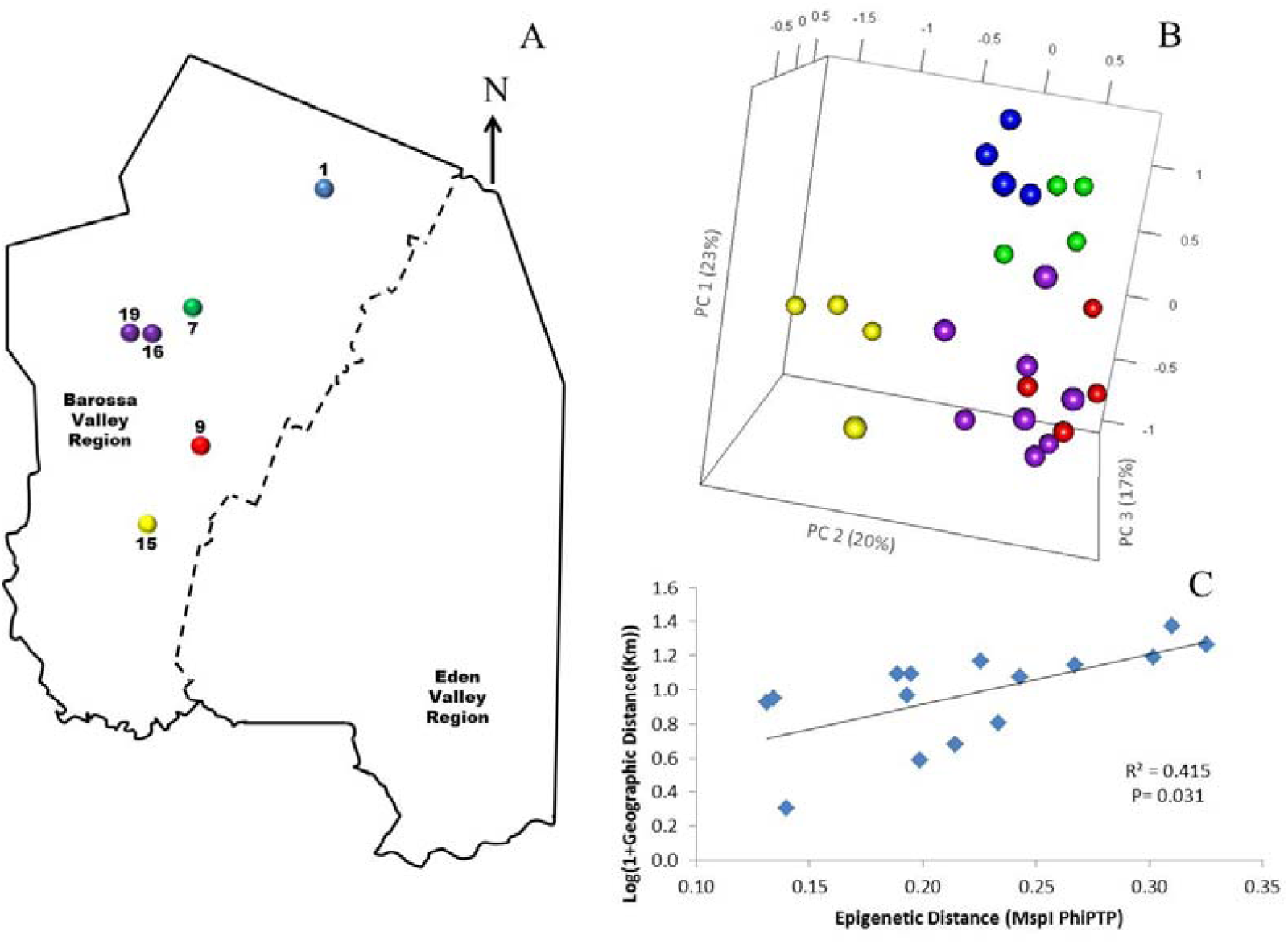
Analysis of the correlation between epigenetic differentiation and geographic distance of vineyards planted with clone 1654 in the Barossa Region and trained using the spur pruned method: **(A)** Location of the selected Barossa Valley vineyards: Northern Grounds (blue), Central Grounds (Green), Eastern Edge (red), Southern Grounds (Yellow) and Western Ridge (Purple). Arrow indicates the direction of geographic North. **(B)** Principal Coordinate Analysis (PCoA) representing epigenetic differences between leaf samples collected from 4 plants/vineyard. Percentage of the variability capture by each Principal Coordinate (PC) is shown in parenthesis. **(C)** Correlation between pairwise epigenetic distance (*Msp*I PhiPT) and geographical distance (Log(1+GeoD (km)) between vineyards. Shown equations are the correlation’s R^2^ and the Mantel test significance (P value was estimated over 9,999 random permutations tests). PCoA and PhiPT for Mantel test are based on presence/absence of 215 loci obtained from MSAP profiles generated using *Msp*I.

Vintage, geographic location and vineyard management have been shown to influence both vegetative growth (Jackson and Lombard 1993) and fruit quality in grapevine (Roullier-Gall et al., 2014). In light of the results shown here, we propose that epigenetic processes in general and DNA methylation in particular, could constitute an important set of molecular mechanisms implicated in the effect that provenance and vintage has, not only on plant vegetative growth, but also on fruit and wine quality. It is also tempting to speculate that in long living crops, such as grapevine, epigenetic priming (Tricker et al., 2013) could allow for the storage of environmental information that would ultimately contribute, at least partially, to the uniqueness of wines produced in different regions.

## Author contributions

HX and MK carried out the experiments and contributed to data analysis. NS performed gene ontology analysis on msGBS data. KGT performed TASSEL analysis on msGBS data. TC, MG, JB, and AM contributed to the design of the research project. RD and CC contributed to the design of the research project, site selection and collection of material. CMRL contributed to the design of the research project, data analysis and drafted the manuscript. All authors read and contributed to the final manuscript.

## Acknowledgements

The authors would like to gratefully acknowledge the Barossa Grounds Project and in particular the growers that allowed us to sample material from their properties and supplied information about their vineyards and management strategies. Dr Kendall R. Corbin performed DNA extractions from all samples used in this study. Personnel in the viticulture group, Dr Sandra Milena Mantilla, Annette James, and Valentin Olek contributed to collection of material.

## Funding statements

This study was funded through a Pilot Program in Genomic applications in Agriculture and Environment Sectors jointly supported by the University of Adelaide and the Australian Genome Research Facility Ltd (Adelaide node). MK was supported by the Australian Agency for International Development (AusAID) PhD scholarship. CMRL is supported by a University of Adelaide Research Fellowship. MG is supported by the Australian Research Council through Centre of Excellence (CE1400008) and Future Fellowship (FT130100709) funding.

---

**Figure S1:**
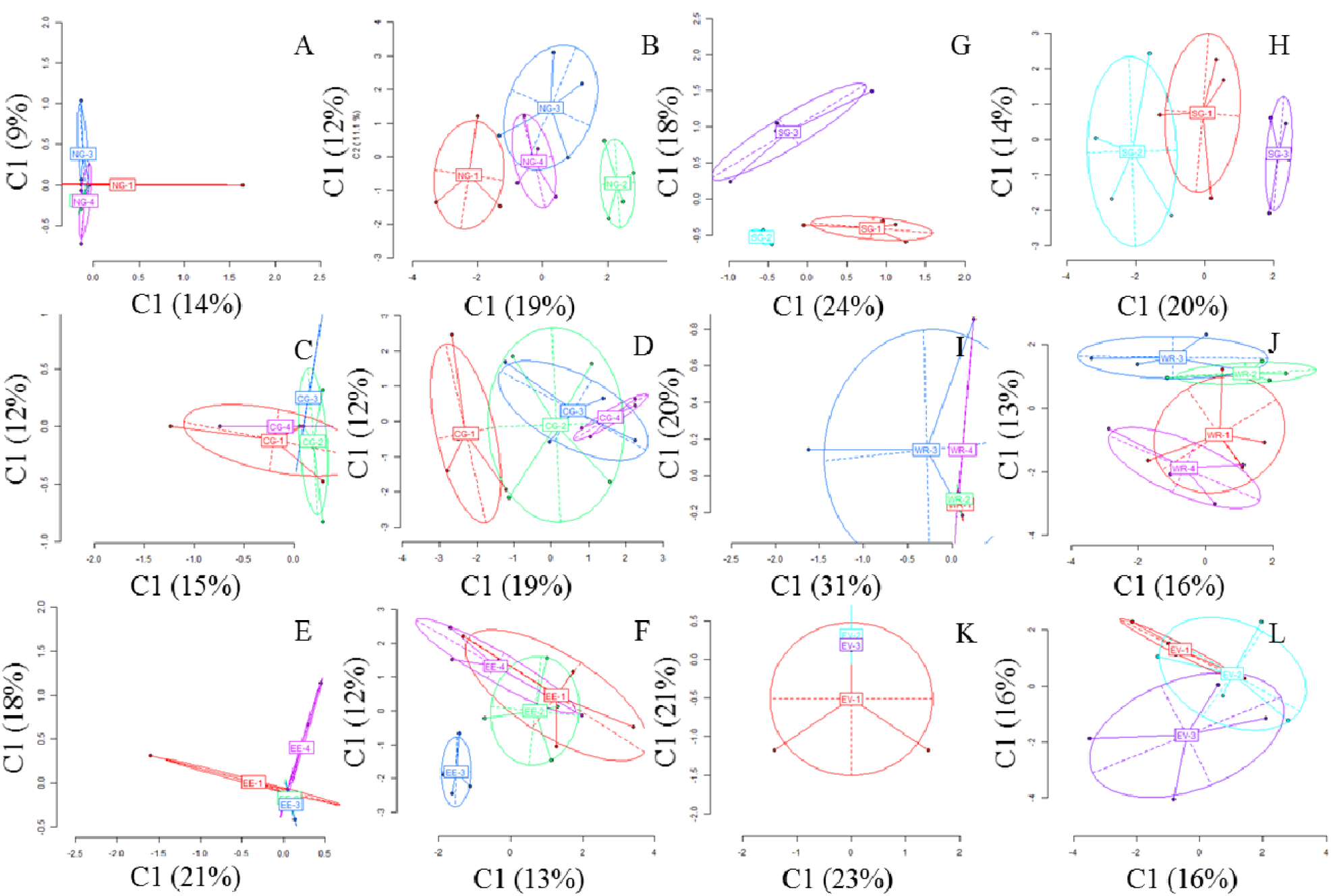
Analysis of genetic and epigenetic differences within Barossa Valley sub-regions: PCoAs represent variability of non-methylated polymorphic loci (Genetic variability) **(A, C, E, G, I and K)** and of methylation sensitive polymorphic loci (Epigenetic variability) **(B, D, F, H J and L)** as classified by the msap (v. 1.1.8) software (Pérez-Figueroa, 2013) of leaf samples in vineyards from Northern Grounds (**A-B),** and the Barossa Valley’s Western Ridge Central Grounds (**C-D)**, Eastern Edge (**E-F)** and Southern Grounds **(G-H),** Western Ridge **(I-J)** and Eden valley **(K-L)**. Coordinates 1 and 2 are shown with the percentage of variance explained by them. Points represent individuals from each vineyard. Vineyard code (NG, CG, EE, SG, WR and EV) is shown as the centroid for each vineyard. Ellipses represent the average dispersion of those poinst around their centre. The long axis of the ellipse shows the direction of maximum dispersion and the short axis, the direction of minimum dispersion.

**Figure S2.**
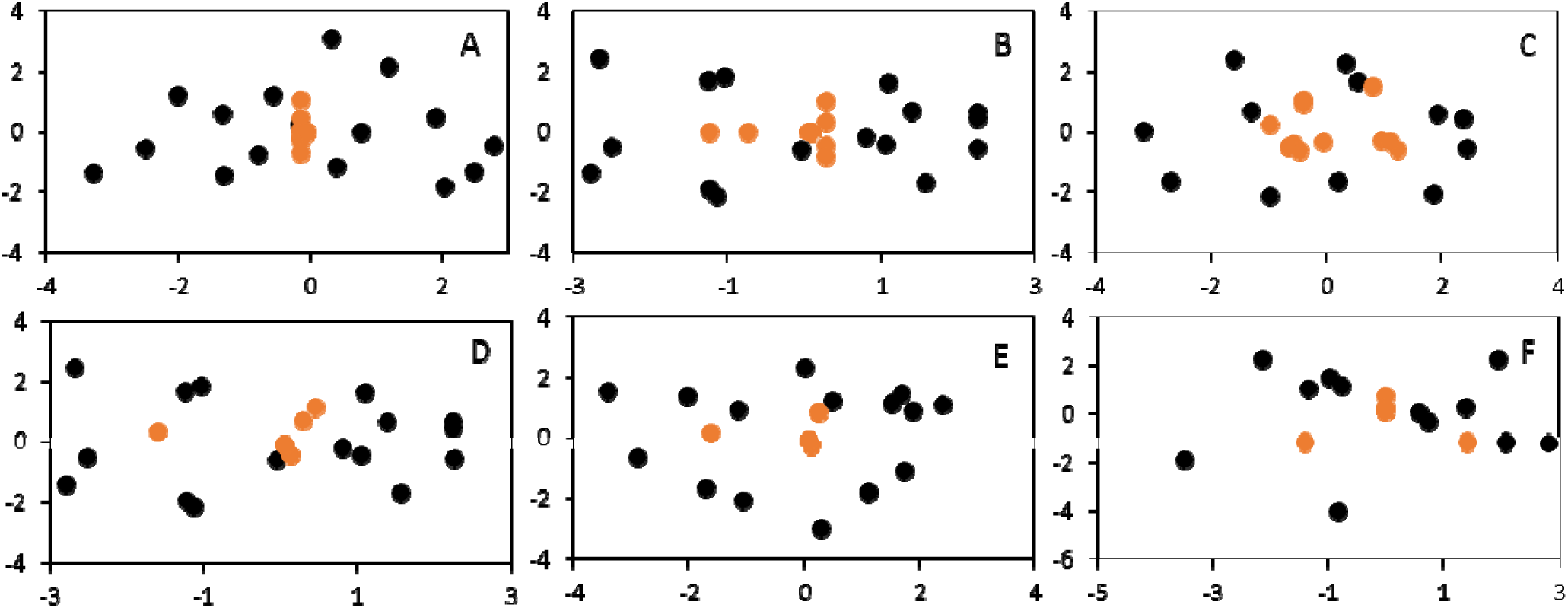
Analysis of regional genetic and epigenetic diversity. Red symbols indicate samples analysed using genetic information only, black symbols represent samples analysed using epigenetic information only according to the R package for Statistical analysis of Methylation-Sensitive Amplification Polymorphism data “msap”. PCoAs were calculated using MSAP profiles generated from gDNA extracted from E-L 7 stage leaves (Coombe, 1995) of 88 grapevine plants grown in vineyards located in the six Barossa Valley wine sub-regions (**A** Northern Grounds, **B** Central Grounds, **C** Southern Grounds, **D** Eastern Edge, **E** Western Ridge, **F** Eden valley) (4 individual plants per vineyard) using restriction enzyme combinations *Msp*I/*EcoR*I and *Hpa*II/*EcoR*I.

**Figure S3.**
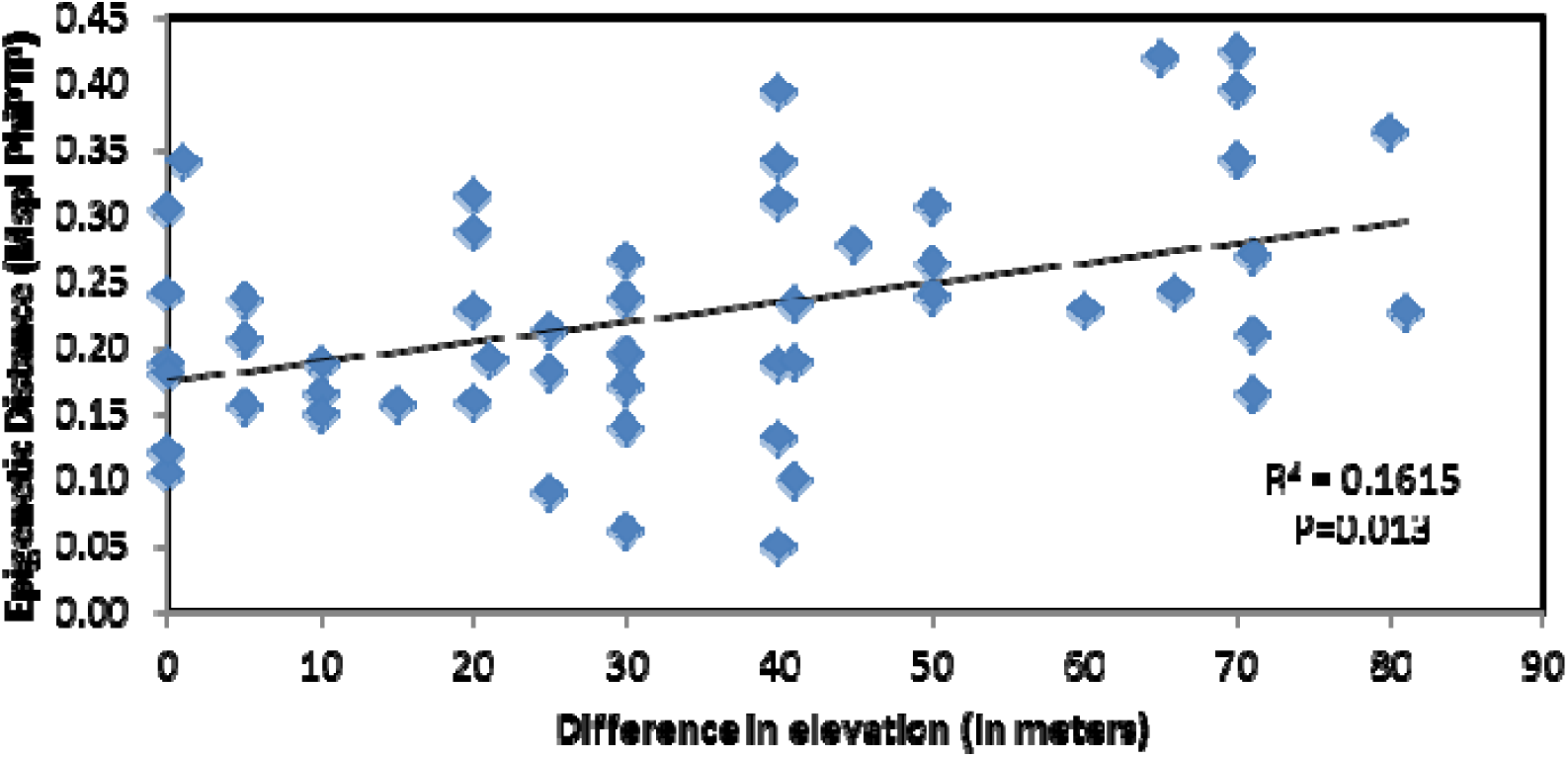
Analysis of the correlation between epigenetic differentiation and environmental differences among vineyards planted along the Barossa Valley North-South axis: Mantel test analysis of the correlation between pairwise epigenetic distance (MspI PhiPT) and differences in altitude between vineyards. Shown equations are the correlation’s R^2^ and the Mantel test significance (P value was estimated over 9,999 random permutations tests). PhiPT values were based on presence/absence of 215 loci obtained from MSAP profiles generated using *Msp*I.

**Figure S4:**
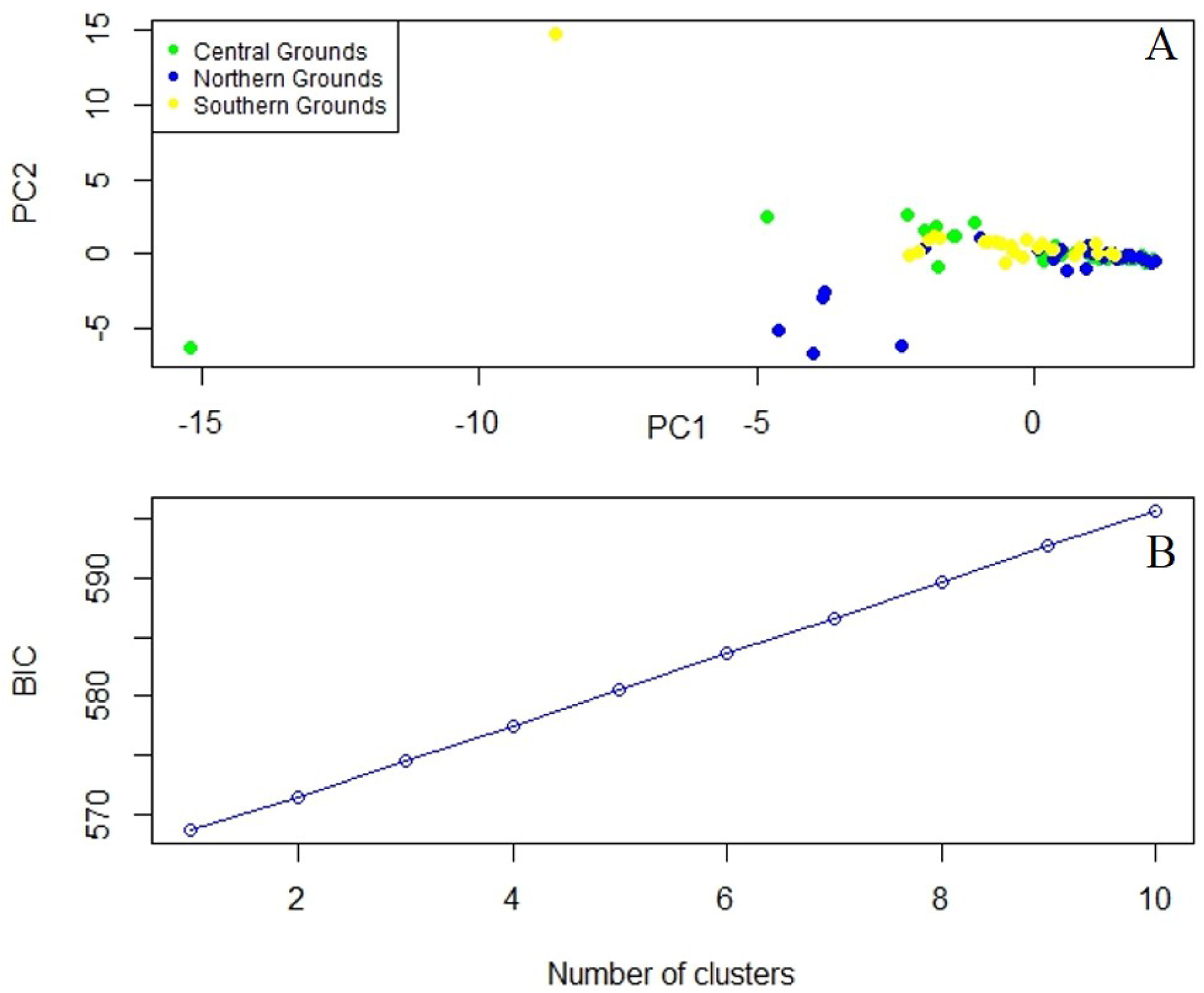
Analysis of the grapevine genetic diversity in vineyards planted along the Barossa Valley North-South axis: **(A)** Principal Component Analysis (PCA) representing genetic structure calculated using 4893 high quality SNPs (i.e. present in at least 80% of the samples) in genomic DNA collected from 11 vineyards (Northern Grounds (blue) 4 vineyards, Central Grounds (Green) 4 vineyards, and Southern Grounds (Yellow) 3 vineyards (9 plants/vineyard)). **(B)** Identification of the optimal number of genetic clusters present within the three sub-regions compared using Bayesian Information Criterion (BIC) as implemented by Discriminant Analysis of Principal Components (DAPC) using adegenet 2.0.0 (i.e. the optimal clustering solution should correspond to the lowest BIC).

**Table S2:**
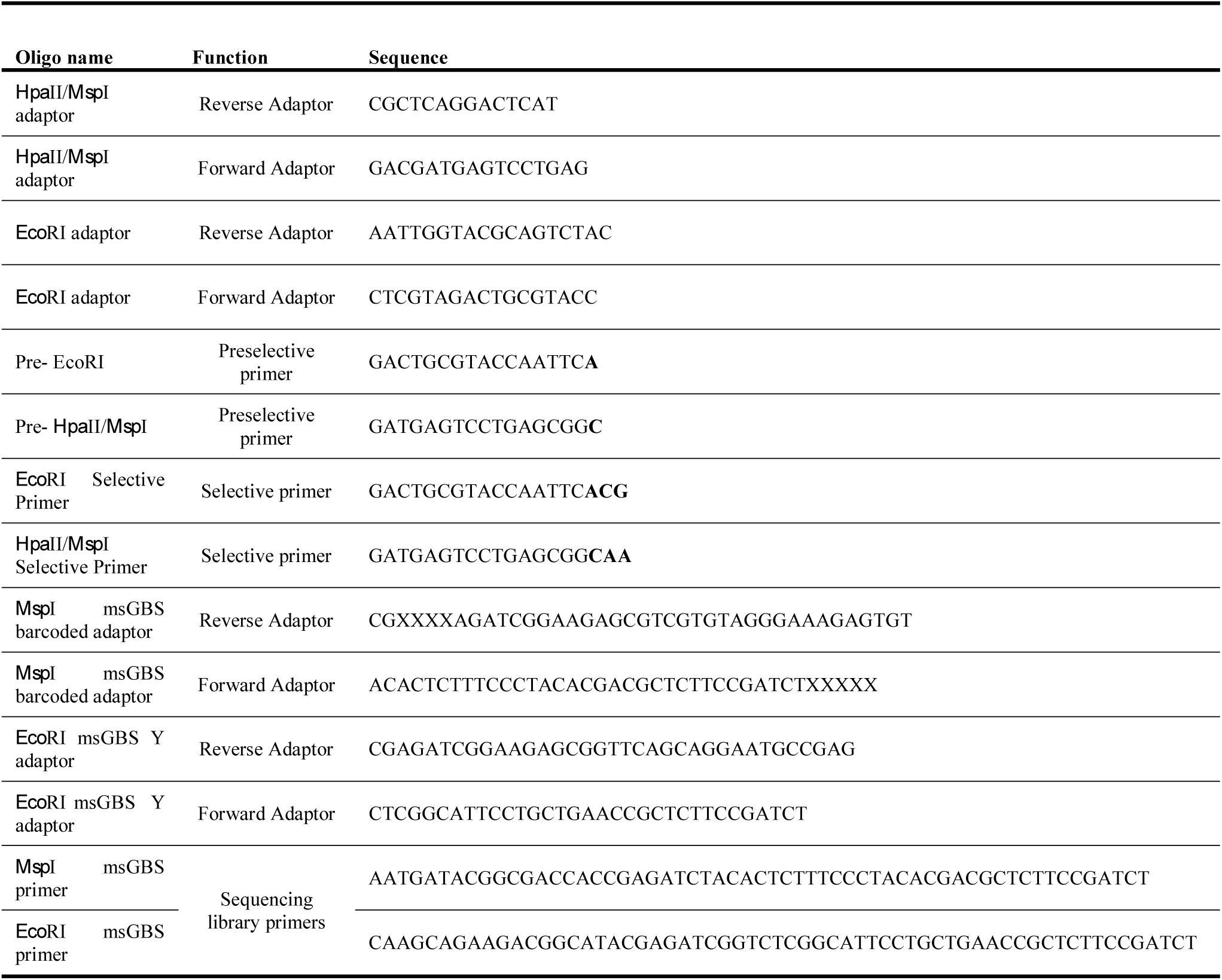
Sequences of oligonucleotide used for MSAP. Selective bases in the primers used during the preselective and selective amplifications are highlighted in bold. Unique msBGS barcode bases are represented as X.

